# Directing Protein Design Choices by Per-Residue Energy Breakdown Analysis with an Interactive Web Application

**DOI:** 10.1101/2023.02.27.530287

**Authors:** Felipe Engelberger, Jonathan D. Zakary, Georg Künze

**Affiliations:** Institute for Drug Discovery, Leipzig University, Brüderstraße 34, 04103 Leipzig, Germany

**Keywords:** Protein design, machine learning, energy calculations, amino acid interactions, web application

## Abstract

Recent developments in machine learning have greatly facilitated the design of proteins with improved properties. However, accurately assessing the contributions of an individual or multiple amino acid mutations to overall protein stability to select the most promising mutants remains a challenge. Knowing the specific types of amino acid interactions that improve energetic stability is crucial for finding favorable combinations of mutations and deciding which mutants to test experimentally. In this work, we present an interactive workflow for assessing the energetic contributions of single and multi-mutant designs of proteins. The **en**ergy break**d**own g**u**ided p**r**otein d**e**sign (ENDURE) workflow includes several key algorithms, including per-residue energy analysis and the sum of interaction energies calculations, which are performed using the Rosetta energy function, as well as a residue depth analysis, which enables tracking the energetic contributions of mutations occurring in different spatial layers of the protein structure. ENDURE is available as a web application that integrates easy-to-read summary reports and interactive visualizations of the automated energy calculations and helps users selecting protein mutants for further experimental characterization. We demonstrate the effectiveness of the tool in identifying the mutations in a designed polyethylene terephthalate (PET)-degrading enzyme that add up to an improved thermodynamic stability. We expect that ENDURE can be a valuable resource for researchers and practitioners working in the field of protein design and optimization. ENDURE is freely available for academic use at: http://endure.kuenzelab.org.

## 1 Introduction

The design of proteins with improved stability and activity is a critical aspect of research in biotechnology and related fields. It holds the potential to revolutionize a wide range of applications (Arnold, 2018), from developing enzymes for industrial processes (Chen and Arnold, 2020), antibodies and antivirals for medicine (Sevy and Meiler, 2014; Willis et al., 2015) to molecular switches and biosensors (Stein and Alexandrov, 2015; Quijano-Rubio et al., 2021). Protein design has been demonstrated to play a crucial role in many different areas of biotechnology (Castro et al., 2022; Habibi et al., 2022; Reetz, 2022).

Two widely used protein design approaches are directed evolution and computer-aided protein design. The former approach mimics the natural gene diversification and selection process and involves iterative rounds of mutagenesis, which create a library of mutants, and selection of mutants with desired functions (Arnold, 2018). Computer-aided protein design typically involves algorithms that suggest mutations for experimental testing (Pan and Kortemme, 2021). These algorithms may be based on in-depth molecular modeling, e.g. with the Rosetta software suite (Leman et al., 2020), or machine learning predictions (Dauparas et al., 2022). The experimental testing of the designed proteins is time-, cost-, and labor-intensive. Thus, prioritizing the most probable candidates for experimental testing is necessary. To facilitate mutant selection, it can be informative to determine the specific types of amino acid interactions that contribute to protein stability and assess the energetic impact of mutations (Goldenzweig et al., 2016).

Machine learning algorithms have revolutionized the field of protein design, enabling researchers to generate novel proteins with improved properties more efficiently (Dauparas et al., 2022; Ferruz and Höcker, 2022). Two current state-of-the-art methods are the evolutionary and structure-based design method PROSS (Goldenzweig et al., 2016; Weinstein et al., 2021), and the deep learning method ProtMPNN (Dauparas et al., 2022). Both methods can yield tens to hundreds of candidate structures or sequences. Selecting the best candidates for experimental testing is a rather tricky task, particularly for designed proteins with multiple mutations, as the effect of each mutation is dependent upon the presence of other mutations – a phenomenon referred to as epistasis (Starr and Thornton, 2016). Yet the selection process determines the success of the overall design process. Thus, it is essential to have reliable, comprehensive, and easy-to-use methods for evaluating and selecting the most probable designs, based on the energetic magnitude and type of interactions (hydrogen bonds, salt bridges, etc.) introduced by the mutations.

Some existing tools for protein design and mutant selection include Rosetta (Leman et al., 2020), HotSpotWizard3.0 (Sumbalova et al., 2018), ProteinSolver (Strokach et al., 2021), and FoldX (Schymkowitz et al., 2005). However, these tools often require extensive knowledge of the software, leaving non-expert users without easy access to analyze the outcomes of their protein design experiments. In the field of *de novo* protein design, DE-STRESS (Stam and Wood, 2021) has been developed to help non-expert users evaluate the plausibility of such designs. Unfortunately, equivalent tools for assessing the results of sequence design experiments operated on a provided structure are lacking. To address this gap, we have developed ENergy breakDown gUided pRotein dEsign (ENDURE), a modular web application that provides an interactive and user-friendly interface for analyzing the energetic contributions of protein designs. ENDURE integrates easy-to-read summary tables and interactive visualizations of automated energy calculations, helping users to explore and reveal mutational hotspots – which confer stabilization or destabilization – and compare the specific types of interactions that a particular mutation is introducing. In that way, ENDURE helps selecting the best protein mutants for further experimental characterization.

The application workflow (**Figure 1**) integrates several key algorithms, which analyze the protein structure using the Rosetta energy function, including per-residue energy breakdown and the sum of interaction energies calculations. Additionally, the tool provides a residue depth analysis, which enables users to track the energetic contributions of mutations occurring at different spatial layers of the protein structure, thus easily shedding light on the particular strategies that a design pipeline might have used and assessing its particular energetic impact. We demonstrate the use of ENDURE in assessing a previously designed version of a polyethylene terephthalate (PET)-hydrolyzing enzyme from *Ideonella sakaiensis* (*Is*PETase), called DuraPETase (Cui et al., 2021), carrying ten mutations compared to the wildtype *Is*PETase (Yoshida et al., 2016). We expect that ENDURE will be a valuable resource for protein designers, filling a crucial gap in assessing and explaining the outcomes of protein design calculations.

**Figure 1.**
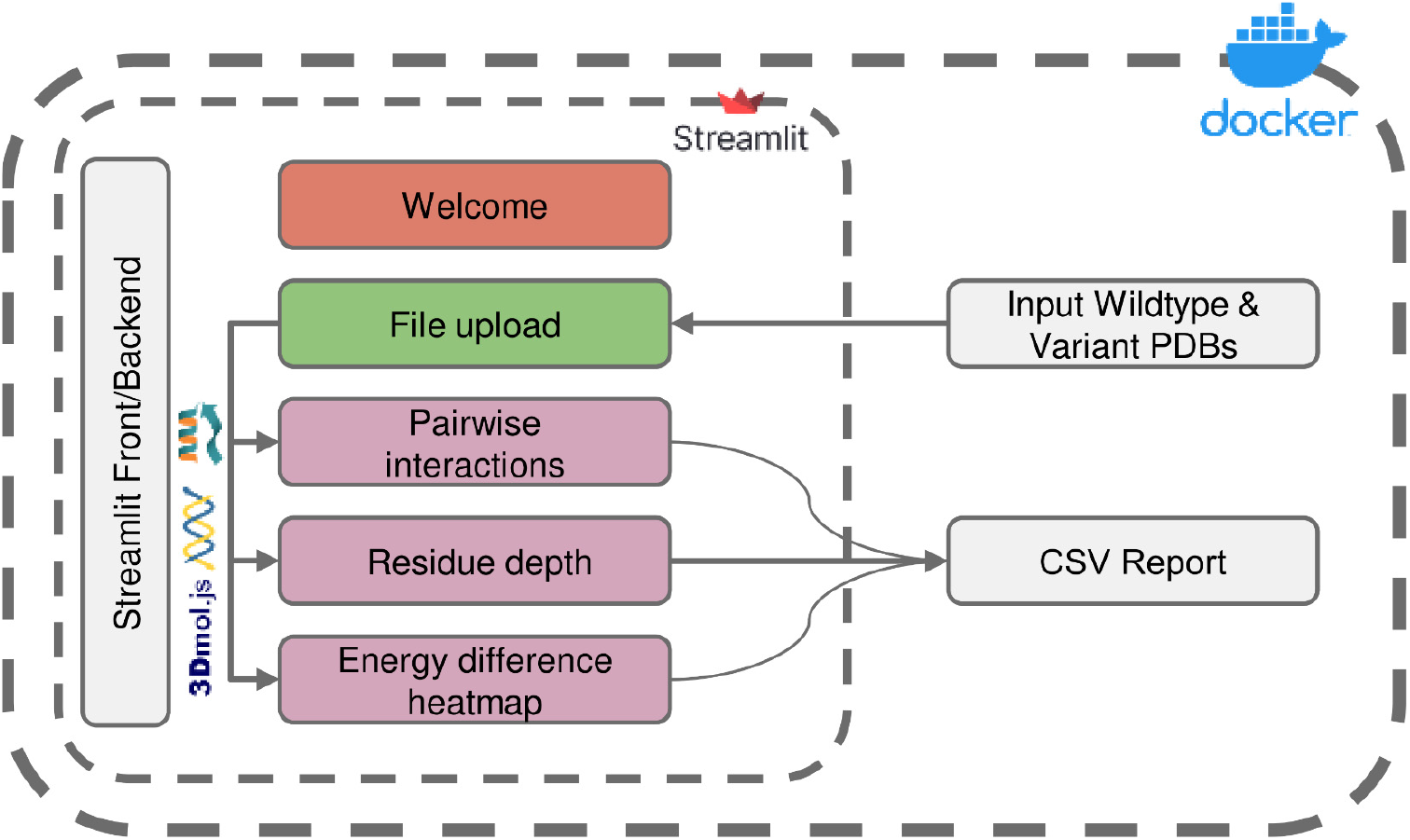
Overview of the ENDURE application architecture. Each colored box represents a subpage that performs a specific task, such as structure pre-processing and Rosetta calculations (File Upload page), pairwise interactions and residue depth analyses, and lastly visualization of an energy difference heatmap. The symbols represent the tools and libraries used for analysis and visualization. The Rosetta software is used for residue interaction analysis, the Biopython library is used for residue depth calculation, and the 3Dmol.js library is used for generating the visualization of the 3D structures. The process starts with uploading the protein structures, and then proceeds to processing and analysis of the structures. Pairwise interactions and residue depth are analyzed, and a CSV report can be generated. Finally, an energy difference heatmap is created.

## 2 Methods

The architecture and interface of the ENDURE web-app are designed in such a way that users are guided through the process of analyzing the energetic contributions of their protein designs in an intuitive and user-friendly manner. As shown in **Figure 1**, the whole workflow consists of several pages implemented in Streamlit (https://streamlit.io), a web application framework for Python. Starting from the File Upload page, the user uploads the PDB files of the designed protein and the reference protein structure. Alternatively, users can choose to provide amino acid sequences of the reference and mutant protein, and ENDURE will automatically predict their structures using ESMFold (Rives et al., 2021). The app then guides the user through three analysis steps, including pairwise interaction analysis, residue depth analysis, and inspection of all pairwise interaction changes by means of an energy difference heatmap. The results of these analyses are presented to the user in an interactive way that can be easily adapted to suit their specific needs using the controls on the left sidebar. The resulting tables can also be exported and downloaded as CSV file. Streamlit abstracts both front-end and back-end programming, making the application easily extensible by users with Python programming knowledge. Finally, the source code of ENDURE is packaged in a Docker container for easy installation and replication of the tool across different compute environments such as servers or HPC clusters. More details on the technological implementation of the app and the computational algorithms are provided in the following paragraphs.

### 2.1 Front End / Backend in Streamlit

The application’s front end consists of the user interface (UI), which includes the welcome page, file upload section, and interactive visualizations. The UI is designed to be user-friendly and easy to navigate, allowing users to upload their protein structure file, prepare the structure, and run the energy breakdown and residue depth calculations. Since Streamlit is our web development framework, the front and backend are managed under the hood. We do include several analysis modules, which are either pure Python functions or Python functions that call other auxiliary executables. The names of important internal functions in the ENDURE app are written in `typewriter font` in the following sections.

### 2.2 Processing structures with ENDURE

The protocol consists of two main stages: (1) structure pre-processing actions and (2) structure analysis actions. Those actions are run on the File Upload page.

The pre-processing is an important step in the ENDURE web tool as it ensures that the uploaded protein structure is in the correct format and ready for analysis. The pre-processing actions include cleaning of the PDB files to remove any ligands, ions, or water molecules, relaxing of the protein structures to remove any steric clashes or unfavorable interactions, and determining the mutations for the designed protein relative to the reference protein. Additionally, the PDB file is renumbered in this step so that the first residue in the file is at position 1. Relaxing the protein structures is the second important action in preparing the protein structures for analysis. This ensures that the protein structure is in a low energy state according to the Rosetta energy function (Alford et al., 2017), which can help to minimize false positive results. The ENDURE web tool uses RosettaScripts (Fleishman et al., 2011) to perform a single iteration of FastRelax (Khatib et al., 2011) by default, taking a previously minimized protein structure and optimizing its energy landscape further. Overall, the input preparation protocol is crucial for ensuring that the protein structure is in the correct format and ready for analysis.

The analysis actions include the energy breakdown (EB) calculations, which provide information about the energetics of each residue’s interaction in the two analyzed protein structures, and the residue depth (RD) calculations. These algorithms are powered by the Rosetta energy function (Alford et al., 2017) and the Biopython (Cock et al., 2009) library, respectively. The EB calculations are performed using the Rosetta EB executable. In short, EB determines the one-body and two-body energies for each residue and decomposes them further into individual score term contributions, thus allowing the simultaneous exploration of, e.g., sidechain and backbone interactions. By clicking on the **Calculate Energy** button on the File Upload page, the Rosetta EB calculation is run in a subprocess. Internally, the `run` function is launched with four parameters as input: the input PDB file name, the location to save the result file, the location to save the log file, and the file path of the Rosetta executable. The output of the protocol is converted to a downloadable CSV file using the `convert_outfile` function, which saves the CSV file as a dictionary in the current session state.

These actions are followed by the RD calculation, which uses the MSMS algorithm (Sanner et al., 1996) from Biopython to calculate the distance of each residue to the molecular surface. The MSMS software computes the solvent-excluded surface from a set of spheres, representing the atoms in a protein structure. The reduced surface is calculated, and an analytical description of the solvent-excluded surface is derived from it. The calculation is done in the `calculate_depth` function.

The processing and analysis actions launched from the File Upload page are run in the background to prevent the UI from freezing, and their results are integrated into the interactive visualizations in the front end.

### 2.3 Analysis of ENDURE outputs: Pairwise interactions analysis

The Interaction Analysis page allows users carrying out a comprehensive analysis of the energetic changes in pairwise interactions of single- and multi-mutant protein designs. The user can select and analyze interactions from different categories (A to F explained below) and from different physical interaction types (salt bridges, etc.) through different control parameters on the side bar. Individual residue pairs, affected directly or indirectly by the mutations, can be selected and displayed in interactive 3D visualizations.

We defined six categories of residue pair interactions to identify the regions in the protein structures most affected by the mutations.

Category A: Residue pairs that are interacting in the reference structure have different interaction energy in the mutant, even though neither of the two residues were mutated.

Category B: Residue pairs that are interacting in the reference structure have one member replaced, resulting in a different interaction energy in the mutant.

Category C: Residue pairs that are interacting in the reference structure no longer interact in the mutant, even though neither of the two residues was mutated.

Category D: Residue pairs that are interacting in the reference structure no longer interact in the mutant because one member was mutated.

Category E: Residue pairs that are not interacting in the reference structure interact in the mutant, even though neither of the two residues was mutated.

Category F: Residue pairs that are not interacting in the reference structure interact in the mutant because one member was mutated.

In order for ENDURE to detect the different interaction categories A-F and different physical interaction types, there are several functions implemented on the page that users can execute by clicking the **Start Calculations** button (after having run all the pre-processing actions on the File Upload page). Such functions perform a post-processing and filtering of the scorefile generated by the Rosetta EB calculations.

The `energy_calc` function is used to identify the essential changes in interaction energies between the mutant and the reference protein structure. It takes the outputs of the residue EB computation performed for the mutant and reference structures, the list of mutations between the reference and mutant, and a streamlit progress bar. The function calls several sub-functions to perform various interaction energy comparisons for each interaction category and physical interaction type. The former is managed by the `interaction_analysis` function and the latter is managed by the following functions: `salt_bridges`, `disulfide_bonds`, and `hydrogen_bonds`. As the names suggest, these functions calculate the energy differences for different types of interactions, such as salt bridges, disulfide bonds, and hydrogen bonds.

The `interaction_analysis` function calculates the difference in interaction energies between a mutant and a reference protein structure for a given list of mutations. The `interaction_analysis` function processes the outputs of the per-residue EB performed on the reference and mutant structures. It calculates the differences in all single-body (i.e., within a single residue) and two-body (i.e., between two residues) energies for all categories (A to F) between the two proteins.

In analyzing the interaction energy changes between the wildtype and mutant protein, it is important to distinguish between total energy changes and significant energy changes. The former is the sum of all energy changes, including those with a small value. However, since thousands of small changes can occur, it is possible for insignificant changes to mask chemically important changes. To overcome this issue, the significant energy change is calculated, which is the sum of only those interaction energy changes that exceed a minimum magnitude. In ENDURE changes that are larger than +1.0 Rosetta Energy Units (REU) or smaller than -1.0 REU are considered significant. By focusing only on significant energy changes, the mutations that are likely to have a significant impact can be more easily detected.

The total energy change for all interactions and the sum of the subset of significant changes, is calculated using the `total_energy_changes` and `significant_changes` functions, respectively. These functions operate on a dictionary returned by `interaction_analysis` and sum all as well as the significant energy changes for each interaction category specified in an interaction list. Interactions from categories A-F can be selected from a list and the change for a given interaction type (salt bridges, disulfide bonds, sidechain-sidechain hydrogen bonds, sidechain-backbone hydrogen bonds, backbone-backbone short-range hydrogen bonds, backbone-backbone long-range hydrogen bonds, and all interactions) can be furthered inspected.

### 2.4 Analysis of ENDURE outputs: Residue depth analysis

Mutation-induced energetic changes can have different effects on different layers of the protein structure. Therefore, we implemented a residue depth analysis and combined it with the per-residue energy breakdown analysis to distinguish changes occurring on the protein surface from those occurring in buried regions of the protein structure.

The Residue Depth page of the ENDURE app allows the user to analyze and compare the effect of mutations on the energy and spatial location of residues. Specifically, the user can select a residue pair that displays a strong negative energetic contribution and visualize the interaction in the protein structure. By adjusting the threshold slider, the user can see which residues have a significant impact on stability and select the most promising mutations for further analysis. Once the user selects a particular mutation by clicking on a point in the scatter plot, the app displays a side-by-side 3D visual comparison of the residue in the mutant and reference structures, which allows for a direct comparison of the effect of mutations on the protein structure. This analysis provides valuable insights into the structural and energetic changes resulting from mutations.

## 3 Results

The Welcome page (**Figure 2**) is the first page users will encounter when accessing the web application. This page provides an overview of the tool and its functionality, allowing new users to quickly familiarize themselves with the interface. The Welcome page includes a brief description of the purpose of ENDURE and the steps involved in using the tool to analyze protein designs. The Welcome page is designed to be user-friendly and intuitive, providing a clear and concise introduction to the tool and its capabilities, making it easier for users to get started and use the tool most effectively for their particular research questions.

**Figure 2.**
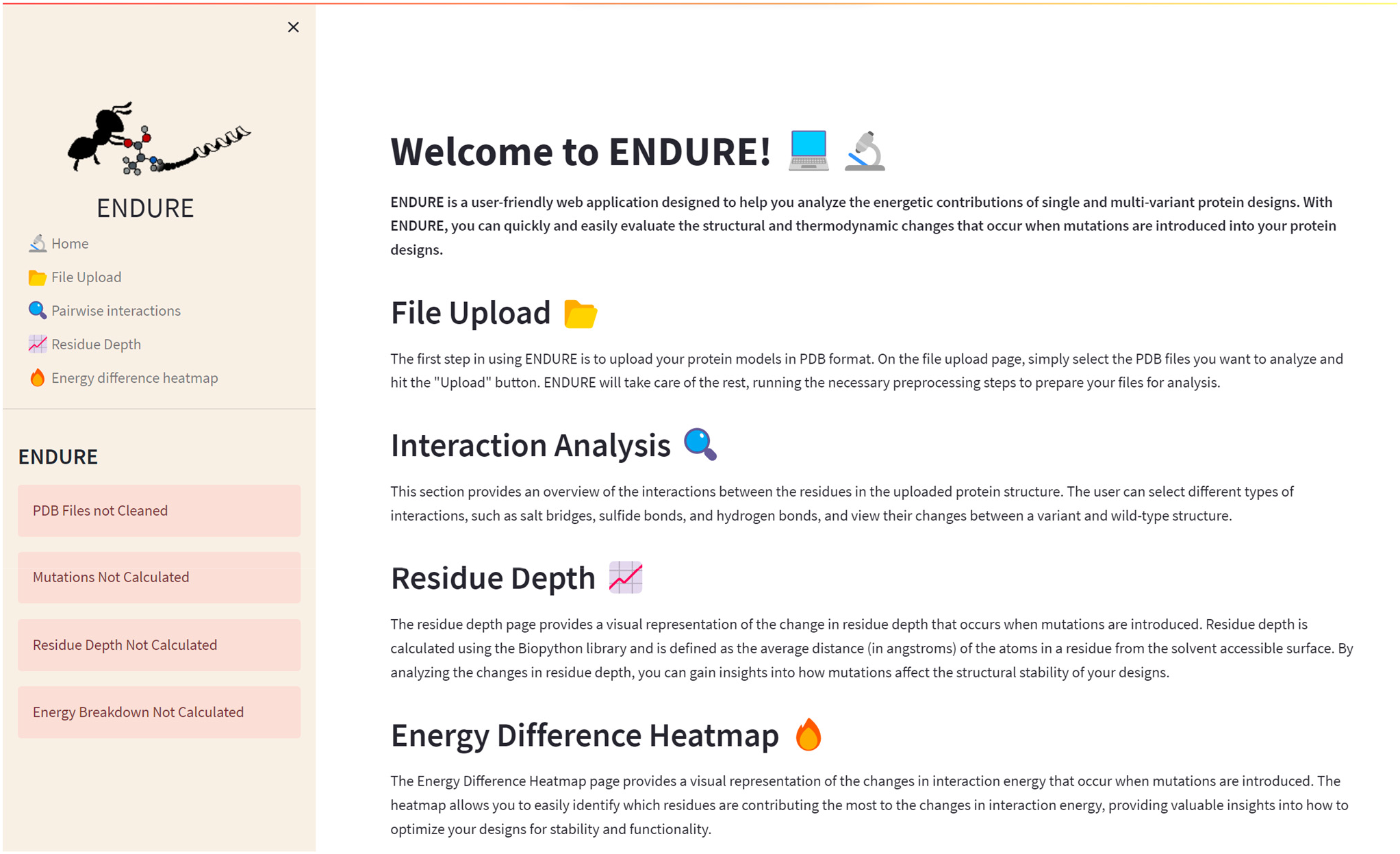
Welcome page. On the left-hand side, the side bar containing the subpages and status bars is given. The latter indicate, by turning from red to green, if the pre-processing and analysis actions on the File Upload page are completed. The main section of the Welcome page contains a brief description of the tool and of each subpage.

Figure 3. presents the File Upload section, where users can upload their protein structure files in PDB format. The interface is designed to be intuitive and user-friendly, with options for either selecting the file from the local file system or dragging and dropping the file into the designated area. After uploading the file, the user is prompted to run five important pre-processing actions: (1) Cleaning PDBs, which ensures that the files are correctly parsed, and the residues are renumbered so that the first residue is at position 1. (2) Relaxing PDB files, which prepares the files for the analysis. (3) Determining mutations by identifying the amino acid differences between the reference and mutant sequences. This information is crucial for many components of the analysis, as it allows tracking the position of mutations. (4) Calculating residue depth determines the average distance of residues from the solvent-accessible surface. This is a key factor in understanding the energetic contributions of mutations. The calculation is performed using the Biopython library and is executed in a separate thread to avoid hanging the GUI. (5) Creating energy breakdown files, which provide a detailed breakdown of the energy contributions of individual residues. This calculation is performed using the Rosetta EB executable, as explained above, and is run in the background to prevent the GUI from being frozen.

**Figure 3.**
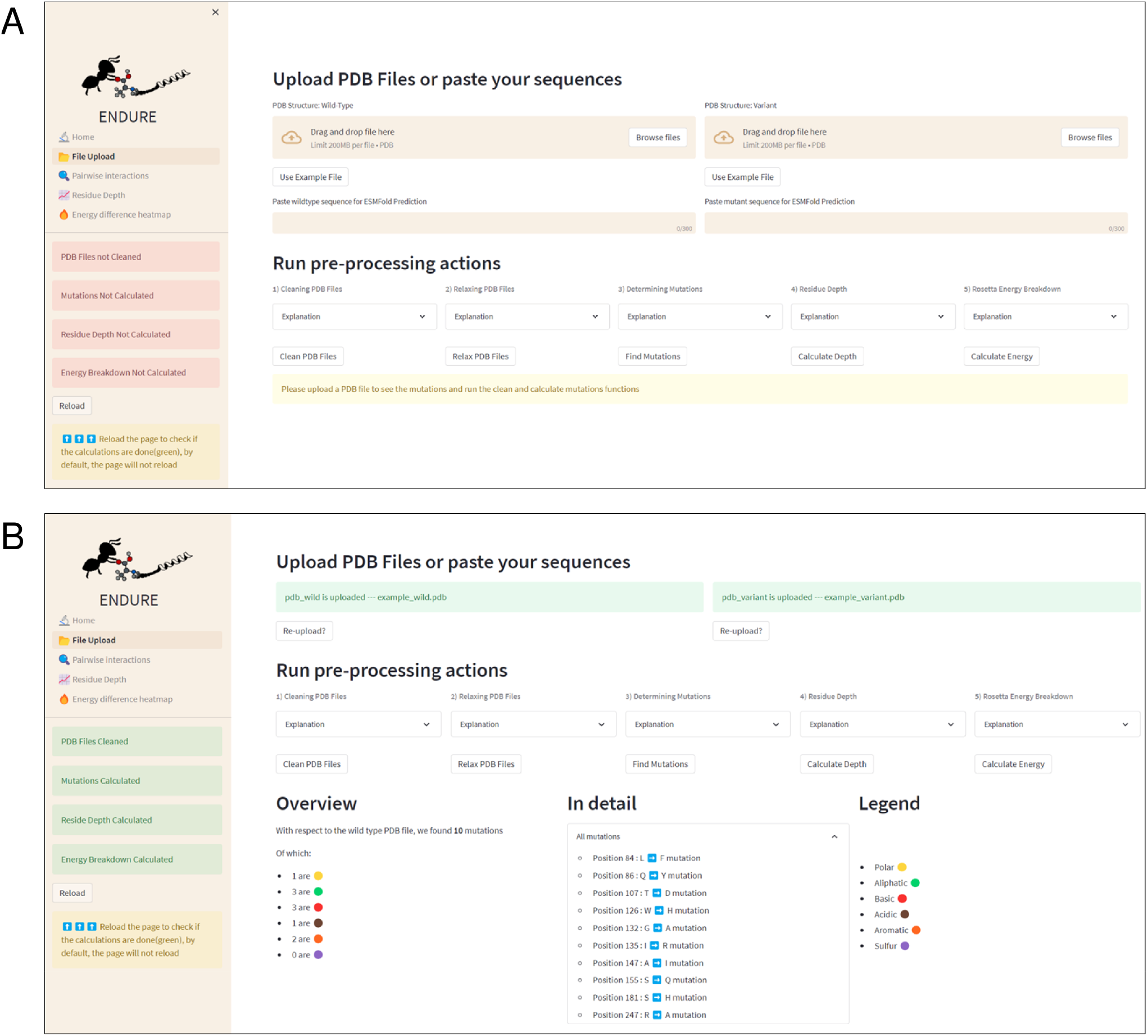
File Upload page. (A) Interface before uploading PDB files or using the example files. The latter action can be activated by clicking the ‘Use Example File’ button. (B) Interface after running all pre-processing and analysis actions. Note that the color of the status boxes turned from red to green.

### 3.1 ENDURE use case: Analysis of an *in silico-designed* PET-degrading enzyme

To demonstrate the workflow and scope of application of ENDURE, we present here the results obtained for a previously reported designed PET-degrading enzyme, called DuraPETase (PDB ID: 6KY5) (Cui et al., 2021), which has higher thermostability than the wildtype *Is*PETase enzyme (PDB ID: 5XJH) (Yoshida et al., 2016; Joo et al., 2018). As can be seen in **Figure 4A**, ENDURE confirms that the particular mutant (carrying ten mutations) has favorable total and significant energy changes (−5.8 REU), indicating improved stability. In addition to the overall energy comparison, ENDURE also provides detailed information about the specific amino acid interactions that contribute to the improved stability of the selected mutant. The interaction analysis feature allows focusing on and visualizing specific residue interactions in the protein structure, which help to rationalize the underlying molecular mechanisms contributing to the improved stability of the selected mutant. These features will be described next.

**Figure 4.**
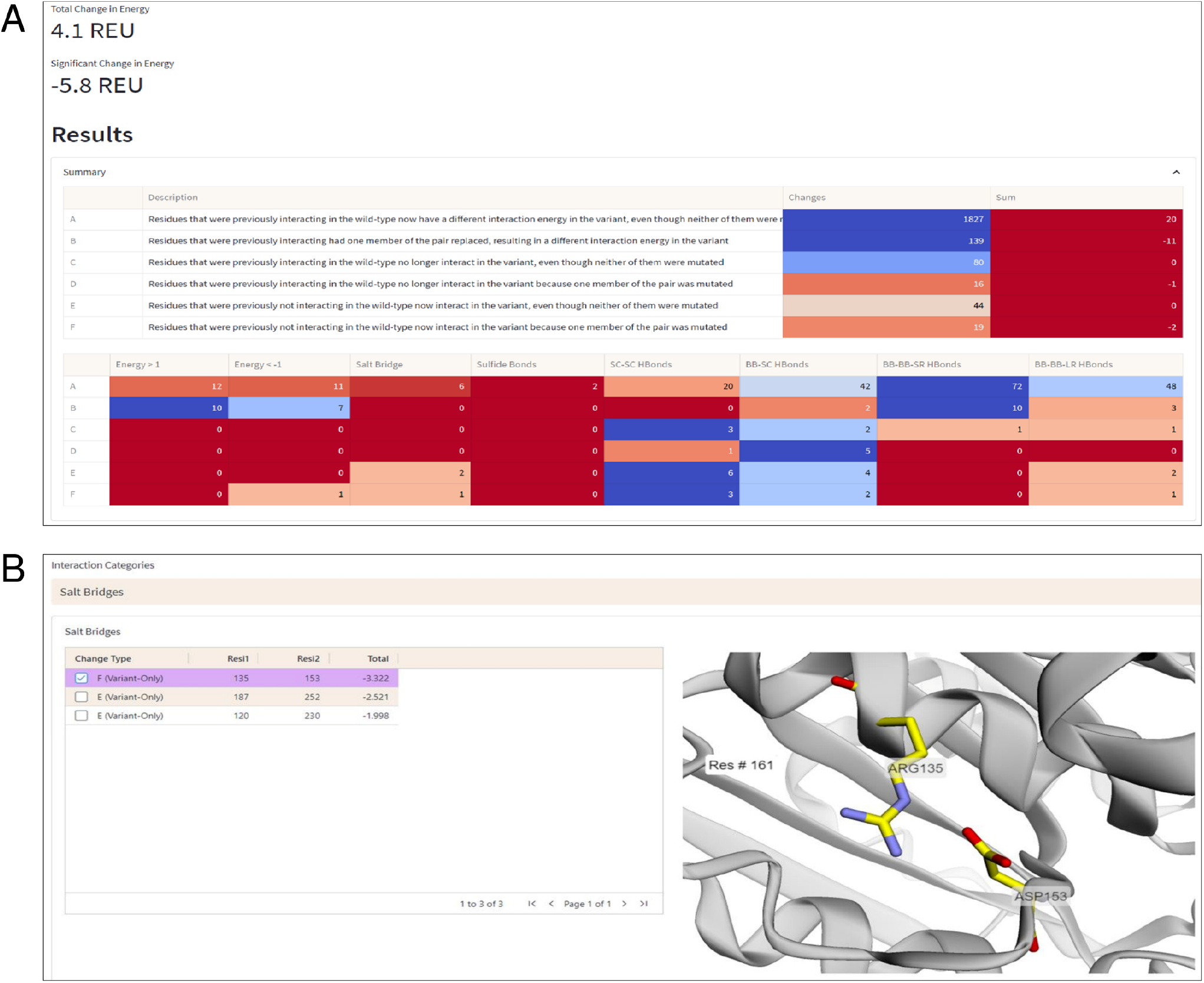
Interaction Analysis page. (A) Summary table of the number of significant energy changes for residue pairs belonging to different interaction categories (A-F, explained in the text) and types of physical interactions in the DuraPETase enzyme. (B) 3D visualization of a salt bridge interaction (Arg135-Asp153) in DuraPETase from interaction category F (i.e., significant energy improvement due to mutation of one residue in the pairwise interaction). Other salt bridges with improved energy in DuraPETase compared to the wildtype *Is*PETase are listed in the table on the left side. The selected Arg135-Asp153 salt bridge is visually represented in the structure viewer window on the right side, highlighting the selected residue pair in sticks. Residue pairs from other physical interaction types (hydrogen bonds between backbone or side chain atoms, disulfide bonds) can also be selected, as explained in the text.

#### 3.1.1 Interaction analysis - Changes in pairwise interactions

The Interaction Analysis page of ENDURE enables performing a detailed examination of the energetic changes for residue pairs of different types (see **Figure 4B**). With this help, the user can identify the particular interactions contributing to improved or impaired energetic stability.

For example, **Figure 4B** shows a salt bridge interaction between residues Arg135 and Asp153, which causes a significant energy improvement of -3.32 Rosetta energy units (REU) in DuraPETase compared to wildtype *Is*PETase. This salt bridge is not present in the wildtype protein but is in DuraPETase – i.e., it belongs to interaction category F. *Is*PETase has an isoleucine (Ile135) at the same position, which cannot form a salt bridge. In addition, as shown in the left-hand table in the screenshot in **Figure 4B**, there are two more salt bridges detected by ENDURE, which contribute significant energy changes in DuraPETase: Asp187-Arg252 (−2.52 REU) and Arg120-Asp230 (−1.99 REU). These two interactions belong to category E, i.e., they do not involve a mutated residue but are probably an indirect effect of nearby mutations in the environment of the four residues. Information like this provides valuable insight into protein structure and can help guide the user in their efforts to design a better-functioning mutant protein.

The different categories of interaction changes (A to F) allow the user to quickly identify the changes in interactions that have occurred and to focus their attention on the most important ones. For example, if a user observes that most changes are of type E or F, this might suggest that new interactions have formed, which could also significantly affect function. By providing this information, the Interaction Analysis page helps the user to quickly understand and prioritize the changes in interactions that have occurred and guide their efforts in protein design.

#### 3.1.2 Residue depth analysis

The residue depth analysis feature allows determining the depth of each residue in the protein structure, which reflects its accessibility to the solvent. By analyzing the energetic changes of mutations occurring in different spatial layers of the protein structure, the location of mutations that improve or impair stability can be determined. In **Figure 5**, the aforementioned mutation Ile135Arg is displayed as an example. In wildtype *Is*PETase, Ile135 is located on the protein surface, indicated by a low residue depth value of ∼2Å (**Figure 5A**). This surface-exposed location is unfavorable for a hydrophobic amino acid. By contrast, the Arg135 residue in DuraPETase can form favorable interactions on the protein surface. In addition to the already mentioned Arg135-Asp153 salt bridge, the side chain of Arg135 interacts with the side chain of Gln155 through a hydrogen bond (**Figure 5B**). The table in **Figure 5B**, which lists the interacting residues for Ile135 or Arg135, respectively, and highlights their respective energy contributions (blue: high energy, red: low energy), confirms these visual observations. These extra interactions, can explain the lower net energy of -14.16 REU for Arg135 in DuraPETase compared to -13.32 REU for Ile135 in *Is*PETase (**Figure 5A**).

**Figure 5.**
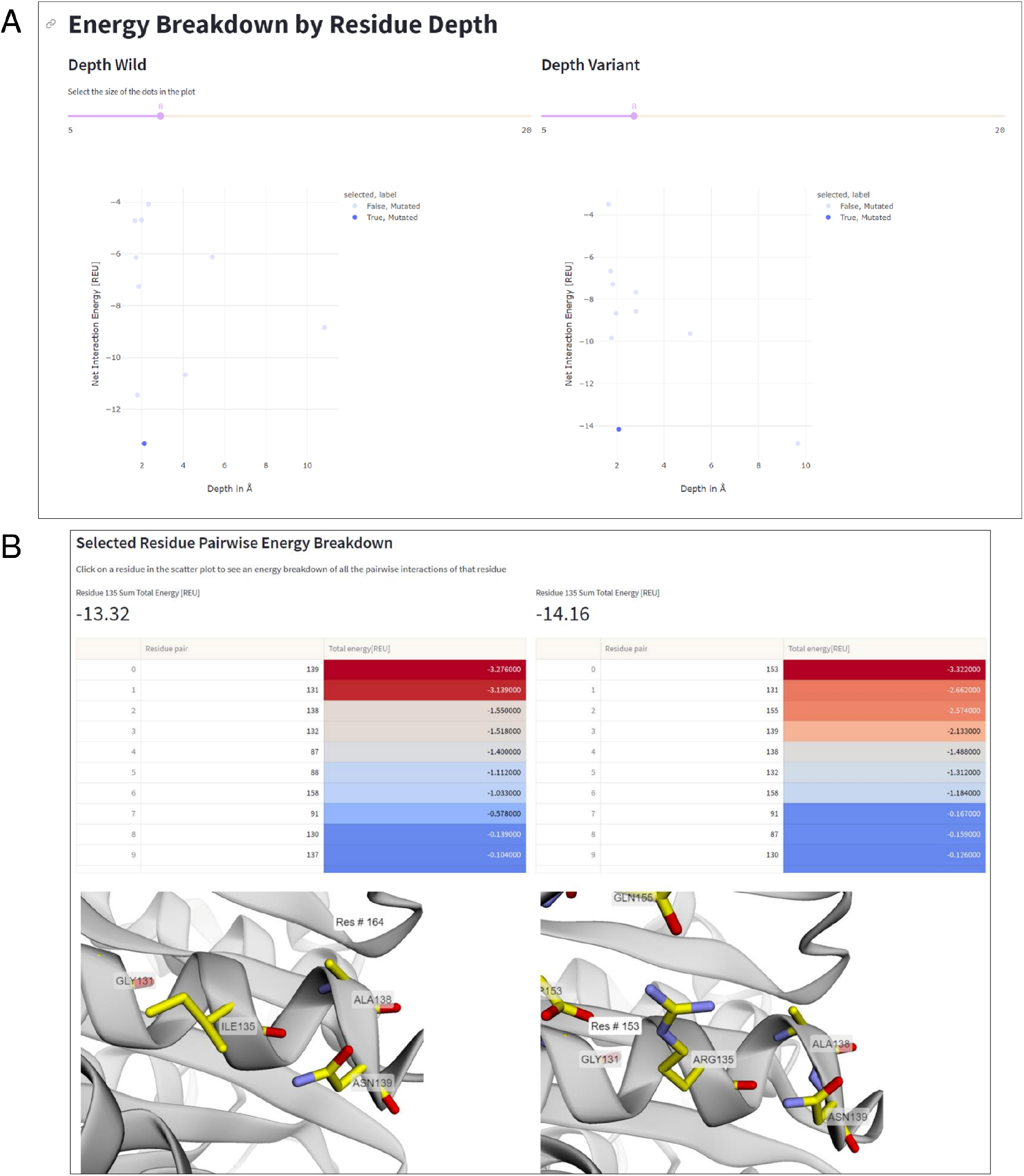
Residue depth analysis. (A) Net interaction energy versus residue depth plots for wildtype *Is*PETA (left) and DuraPETase (right). The blue data point in the lower left corner of the plot, corresponding to Ile135 in *Is*PETase or Arg135 in DuraPETase, respectively, has been selected. (B) Side-by-side comparison of the location and surrounding amino acids of Ile135 and Arg135, respectively. The interacting residues and their energy contributions are listed above the 3D viewer.

Information like this is important because it can help decide which residues to mutate in order to improve protein stability. For example, mutations in the protein core may have a greater impact on stability than those located near the surface, and thus targeting core residues for mutagenesis can considerable impact the stability of the designed protein. Additionally, the residue depth analysis feature can be used to select the most promising mutants for further characterization, by identifying mutations that have an improved net interaction energy and are located in favorable positions within the protein structure.

#### 3.1.3 Energy difference heatmap

This feature allows the user to quickly represent all residue pairs that have a significantly changed interaction energy in the mutant protein compared to the wildtype, and selectively track specific pair interactions. The user can select interactively from the residue pair interaction matrix on the left side in **Figure 6** a pair of residues that display a large negative energetic contribution. The relevance threshold for the energy can be adjusted with the threshold slider present above. Once the user has selected a residue pair (**Figure 6A**), the corresponding pair (Arg135-Asp153 in this example) will be highlighted in the structure viewer in the middle of the page (**Figure 6B**), and a breakdown of the interaction energy change into individual score terms from the Rosetta energy function will appear (**Figure 6C**). For visual clarity only none-zero score terms are displayed to the user.

**Figure 6.**
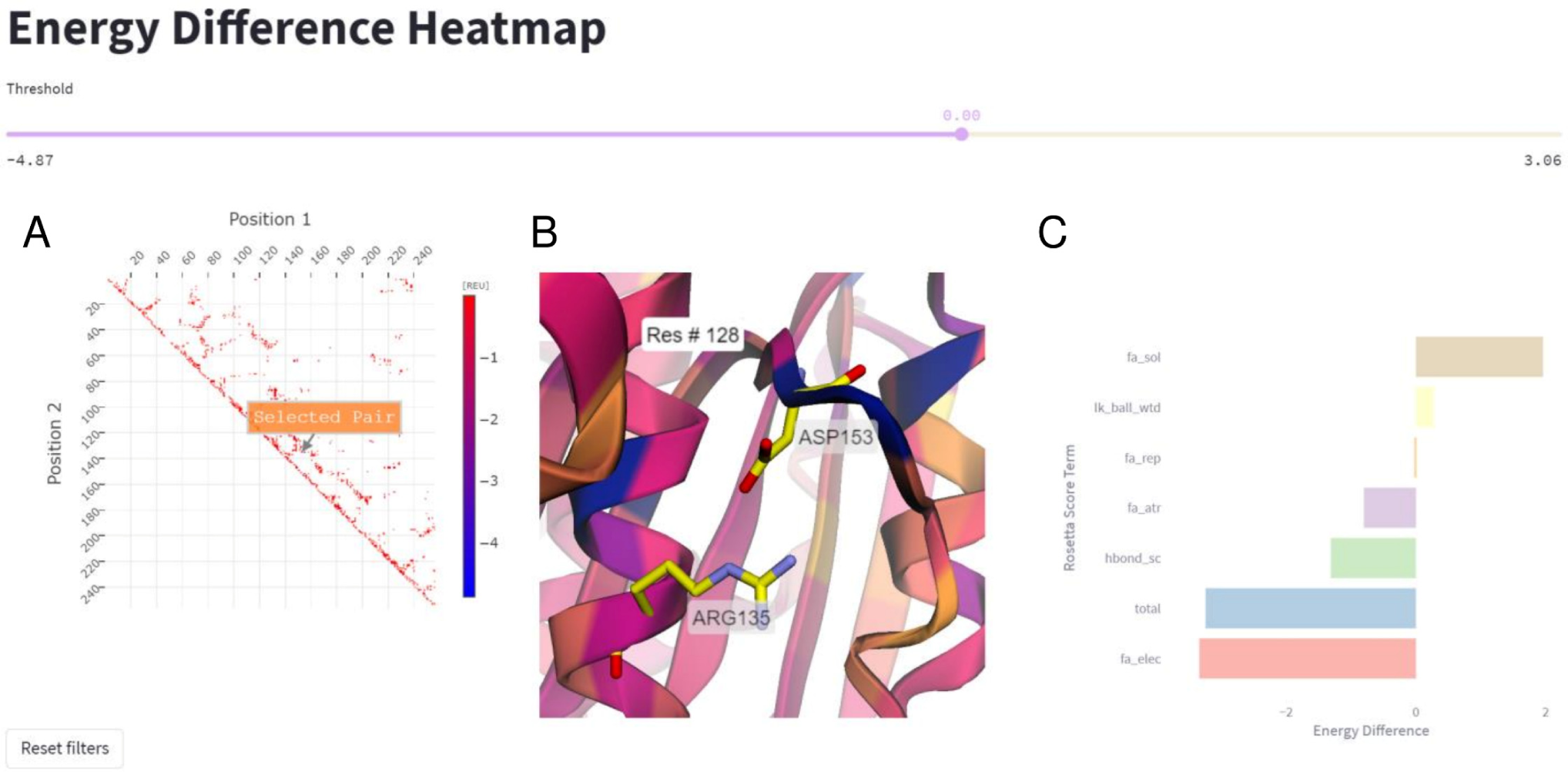
Energy difference heatmap. (A) 2D matrix of residue pair energy changes between *Is*PETase and DuraPETase. Negative values indicate that an interaction has a lower (more negative) energy in DuraPETase. The 2D matrix is illustrated as a heatmap with larger negative changes colored blue and smaller negative energy changes colored red. Users can zoom in the 2D matrix and interactively select a residue pair for further analysis. (B) 3D visualization of the Arg135-Asp153 residue pair after selecting it in the heatmap in A. The structure is colored according to the positional energy difference. (C) Breakdown of the energy change for the selected pairwise interaction into individual Rosetta score terms, representing different physical interactions.

## 4 Discussion

The ENDURE web application provides a user-friendly interface for analyzing protein structures to facilitate mutant selection in protein design workflows. The study has shown that the application can accurately and efficiently process PDB files, clean and renumber them, determine mutations, calculate residue depth, and generate energy breakdown files. The interaction analysis section allows users to view changes in pairwise interactions between residues in the wildtype and mutant structures by providing visual representations of the significant and total energy changes.

One of the key innovations of the ENDURE web application is its ability to group changes in residue pairwise interactions into different types and categories, ranging from residues that are interacting in the reference and mutant protein structure with different interaction energies (category A), to residues that are not interacting in the reference structure but make interactions in the mutant due to a mutation (category F). These different categories of changes provide a useful way for the user to identify which mutations have the strongest impact on the protein structure, and consequently focus design efforts on specific areas of the protein structure.

Compared to previous research, our web application offers a user-friendly and accessible solution for analyzing protein structures and interaction changes. Prior research in this field has typically been focused on developing computational tools for protein structure prediction and analysis (Schymkowitz et al., 2005; Leman et al., 2020; Stam and Wood, 2021). In contrast, our web application provides a unique solution by incorporating these tools into a user-friendly interface and making them more broadly accessible.

It is important to note that the tool is limited by the accuracy of the underlying computational tools and algorithms used for protein structure prediction and analysis. Additionally, the interaction analysis section is based on a static protein structure analysis. Accuracy in predicting mutation-induced energy changes could benefit from a model ensemble approach (Peccati et al., 2023), in which protein dynamic changes like loop rearrangements can be considered. These limitations should be taken into consideration when interpreting the results of the analysis.

Despite these limitations, it represents a significant advancement in the field of protein structure and interaction analysis. The integration of computational tools into a user-friendly interface makes it possible for scientists outside the field of computational structural biology to quickly and efficiently analyze protein structures and identify potential areas for improvement. In future versions, ENDURE could be expanded to include additional features and improvements, such as the ability to examine other kinds of proteins, including membrane proteins and proteins with noncanonical amino acid modifications, through the incorporation of different energy functions. It could also be enhanced to consider ligand molecules in the design analysis, allowing the identification of designs with improved binding affinity. These features are currently planned for the next version of ENDURE, which will be released in the future. Additionally, the tool could be further developed to incorporate machine learning techniques to improve the accuracy of the analysis. With these advancements, ENDURE could become an even more powerful tool for protein design and analysis.

In conclusion, the ENDURE web application provides a unique and accessible solution for analyzing protein structure and interaction changes and, in that way, represents a significant advancement for the field of protein design. Categorizing changes in pairwise interactions for different interaction types provides a straightforward way for the user to guide their protein design strategies. Integrating computational tools into a user-friendly interface makes it possible for a broader audience to quickly and efficiently analyze protein structures. The future direction of the research will focus on further developing the application to incorporate analysis on protein dynamics, support for non-standard amino acid residues, and application of machine learning techniques. Furthermore, a command line interface integrated in the front end is planned, which will help further customize some of the analyses.

## Data availability statement

The source code for ENDURE and example data used as use case in this paper can be downloaded from: https://github.com/kuenzelab/ENDURE. The ENDURE web application is freely available for academic use at http://endure.kuenzelab.org.

## Author contributions

Conceptualization, F.E. and D.Z.; methodology, F.E. and D.Z.; software, F.E. and D.Z.; investigation, F.E. and D.Z.; data curation, F.E.; writing—original draft preparation, F.E.; writing—review and editing, F.E., D.Z. and G.K.; visualization, F.E.; supervision, G.K.; project administration, G.K. All authors have read and agreed to the published version of the manuscript.

## Acknowledgments

D.Z. acknowledges the Max Kade foundation for receiving a summer exchange scholarship. The authors thank Ivan Ivanikov for his technical support in installing the ENDURE webserver. The authors further acknowledge the Leipzig University Scientific Computing Center and the Institute for Drug Discovery for providing the computational resources.

